# Genomic determinants of speciation

**DOI:** 10.1101/837666

**Authors:** Qian Cong, Jing Zhang, Nick V. Grishin

## Abstract

Studies of life rely on classifying organisms into species. However, since Darwin, there is no agreement about how to separate species from varieties. Contrary to a frequent belief, quantitative standards for species delineation are lacking and overdue, and debates about species delimitation create obstacles for conservation biology, agriculture, legislation, and education. To tackle this key biological question, we have chosen butterflies as model organisms. We sequenced and analyzed transcriptomes of 188 specimens representing pairs of close but clearly distinct species, populations, and taxa that are debated among experts. We find that species are robustly separated from populations by the combination of two measures computed on Z-linked genes: fixation index that detects hiatus between species, and the extent of gene flow that quantifies reproductive isolation. These criteria suggest that all 9 butterfly pairs that caused experts’ disagreement are distinct species, not populations. When applied to *Homo*, our criteria agree that all modern humans are the same species distinct from Neanderthals, suggesting relevance of this study beyond butterflies. Furthermore, we found that proteins involved in interactions with DNA (including proteins encoded by *trans-*regulatory elements), circadian clock, pheromone sensing, development, and immune response recurrently correlate with speciation. A significant fraction of these proteins is encoded by the Z-chromosome, which appears to be resistant to introgression. Taken together, we find common speciation mechanisms in butterflies, reveal the central role of Z-chromosome in speciation, and suggest quantitative criteria for species delimitation using genomic data, which is vital for the exploration of biodiversity.

## INTRODUCTION

“I was much struck how entirely vague and arbitrary is the distinction between species and varieties” wrote Darwin in his classic book “On the Origin of Species” (Darwin 1859). Traditionally, biological species are defined by the reproductive barrier: interspecies hybrids are expected to be either sterile or less fit than individuals of either species. However, modern genomic studies of animals (and human) revealed profound gene flow (introgression) between distinct species, blurring the boundaries between species. This relatively new concept suggests that the reproductive barrier between species, especially sister species, is frequently incomplete (Fontaine et al. 2015; Heliconius Genome Consortium 2012; Villanea and Schraiber 2019). The perceived distinctness between animals recognized as different species, despite continuing gene flow between them (Mallet et al. 2016), dazzled many biologists. There is hardly a question as contentious in biology as the definition of species, conceptually and practically (Freudenstein et al. 2017). While the general consensus is that a certain level of reproductive isolation indicates speciation, quantification of such isolation is incomplete. Much attention has been paid to speciation in model organisms, and a number of genes involved in reproductive isolation between species have been identified (Maheshwari and Barbash 2011; Phadnis et al. 2015). With the advent of genomic sequencing, we are poised to advance the understanding of the genetic determinants of speciation and the molecular mechanisms of reproductive isolation in a broader spectrum of organisms, and to define species on the basis of their genotypes.

Introduced by Remington (Remington 1968), a suture zone is a limited geographic area that serves as a common boundary between many narrowly sympatric and frequently very close pairs of species on either side of the suture zone. Most suture zones in North America lie in the western part of the continent and are associated with mountain ranges separating populations (Remington 1968). More interesting and subtle zones are to the east. They lack obvious geographic barriers and offer challenges for finding the isolation mechanisms between species. The central Texas suture zone where western and eastern populations meet has been well-characterized in birds (Newton 2003). Over 20 pairs of bird species are known to be split into western and eastern forms with the boundary in Texas. Remington suggested that the central Texas suture zone was a “mature zone” because many western-eastern species pairs are well isolated reproductively and rarely hybridize when they meet in Texas (Rising 1983).

Extensive characterization of suture zones in butterflies is still lacking. Casual observations reveal several known pairs of butterfly species segregated in western (more precisely, southwestern) and eastern (northeastern) forms around the central Texas suture zone. Several pairs have been studied at the genomic scale, for instance, *Calycopis cecrops* (eastern) and *C. isobeon* (western) (Cong et al. 2017; Cong et al. 2016a), *Phoebis sennae eubule* (eastern) and *P. sennae marcellina* (western) (Cong et al. 2016b), and *Heraclides cresphontes* (eastern) and *H. rumiko* (western) (Cong and Grishin 2018). These studies reveal differences in the degree of genomic separation and hint at some common mechanisms of speciation. To better understand the genomic determinants of speciation across the central Texas suture zone and to probe possible evolutionary scenarios of speciation, we selected 25 pairs (Fig. 1) of butterflies distributed in both eastern and western sides of this suture zone and obtained their RNAseq data. The “eastern” populations correspond to taxa widely distributed over the eastern United States from Texas to Florida and northwards to Canada in many species. The eastern US zone is characterized by colder climate and higher humidity. The “western” populations include taxa that are distributed from central Texas southwards into Mexico and frequently westwards to variable extent, sometimes all the way to the Pacific coast. The southwestern zone is warmer and typically dryer, especially westwards.

**Fig. 1.**
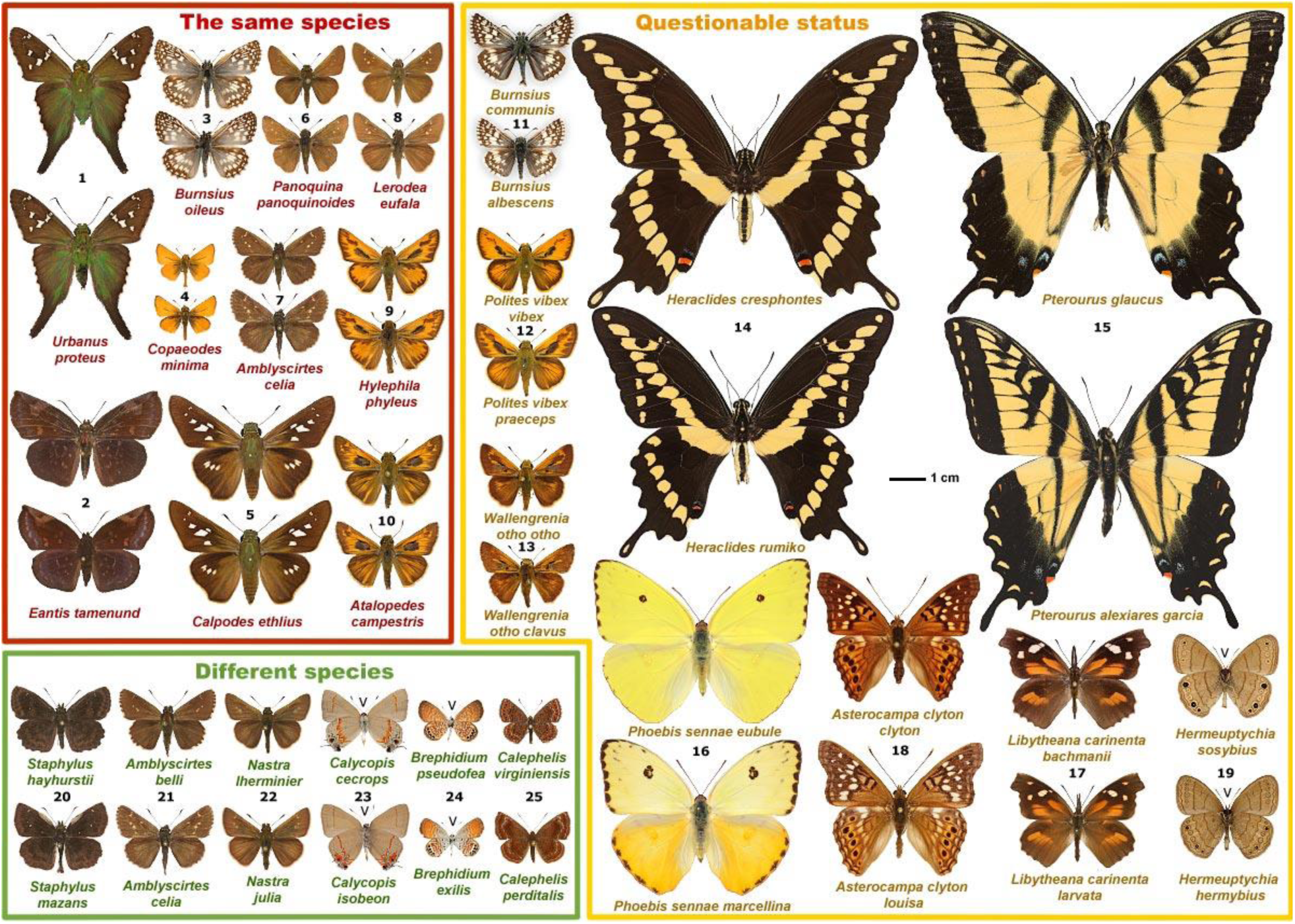
Pairs of taxa around the central Texas suture zone analyzed in this study. Pairs in the red box are expected to be the same species (negative control); pairs in the green box are well-separated, different species (positive control); pairs in the yellow box include possible species or subspecies, but experts disagree on their status (test set). Specimens are shown in pairs of counterparts: eastern is above and western is below, and species names are given below specimens. Dorsal side of each specimen is shown except when marked with “V” between the antennae, which indicates that ventral surface is sown. Numbers shown between the specimens on each pair correspond to those used in Fig. 2.

The 25 pairs selected included (1) positive controls: well-differentiated species of unquestionable distinctness, such as *Nastra lherminier* (eastern) and *N. julia* (western) or *Staphylus hayhurstii* (eastern) and *S. mazans* (western); (2) pairs that may be different species, but opinions differ regarding their taxonomic status: some of these taxa are currently treated as subspecies, i.e., geographic races that differ morphologically but are not expected to show reproductive isolation strong enough to qualify as species, e.g., *Wallengrenia otho otho* (eastern), and *W. otho clavus* (western); and (3) negative controls: species that are known to be morphologically uniform across their distribution from east to west and are not expected to speciate across the suture zone, such as *Lerodea eufala* and *Hylephila phyleus*. By measuring the divergence between eastern and western populations, we aim at detecting general trends of speciation and isolation (or lack of them), and establishing some standard measures for genome-level isolation in the process of speciation.

## RESULTS AND DISCUSSION

### Genomic measures to quantify the reproductive barrier and define species

In practice, species are defined by the hiatus between them that results from at least partial reproductive barriers. Historically, decisions about species boundaries were mostly based on morphology, and animals were partitioned into phenotypically distinct groups separated from each other by discontinuity in variation. Moving to genotypes, a standard measure of genomic distinctness is fixation index (*F*_*ST*_: 1 − *DIV*_*within*_/*DIV*_*between*_, where *DIV*_*within*_ and *DIV*_*between*_ are the divergences within and between populations, respectively). We calculated *FST* over all protein sequences encoded by our sequenced transcripts, and the result is shown in the upper right corner of Fig. 2A (x-axis only). Although *F*_*ST*_ for positive control pairs (well-established different species) are always higher than the negative controls (conspecific pairs), the values of *F*_*ST*_ over all proteins show a continuous distribution without a break between pairs of species and pairs of conspecific populations. To quantify their reproductive isolation, we directly measured the extent of gene flow (*E*_*GF*_) between each pair. We consider an allele to result from gene flow if it is more similar in sequence to alleles from individuals of another group than to alleles from its own group. This measure (y-axis in upper right corner of Fig. 2A) seems to separate the positive from negative controls with a larger gap, but nevertheless forms a continuous spectrum for all analyzed pairs, similar to *F*_*ST*_.

**Fig. 2.**
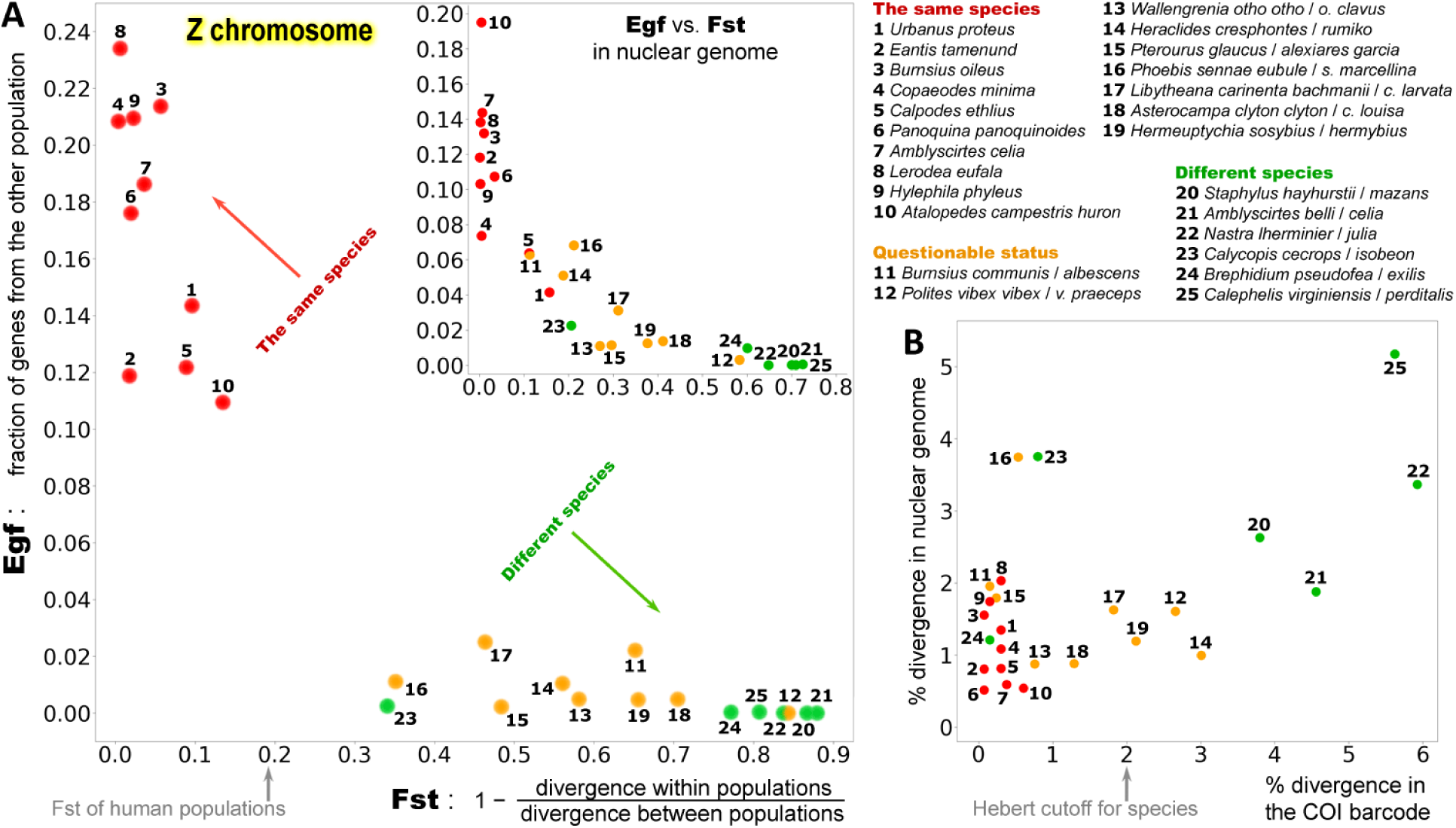
Quantification of the reproductive barrier and genomic criteria for defining species. (A) Fixation index (*F*_*ST*_) and extent of gene flow (*E*_*GF*_) calculated on Z-linked genes clearly distinguish pairs of different species from conspecific pairs, while the same measurements computed on all nuclear genes present in RNAseq (upper right corner) dataset do not separate them as well. (B) Divergence (percent of different nucleotides) in COI barcode and nuclear genes performs poorly in evaluating whether two taxa are different species.

We reason that the continuity in the distribution of *F*_*ST*_ may be caused by incidental hybridization, gene flow and introgression between species, all of which make different species more similar to each other. For example, introgression between *Heliconius* species plays an important role in mimicry, with essential regions regulating wing pattern development being transferred between species (Heliconius Genome Consortium 2012). We also documented introgression of mitochondria between a pair of well-established species used in this study: *Calycopis cecrops* and *C. isobeon* (Cong et al. 2017; Cong et al. 2016a). Because a lower level of introgression in the Z-chromosome was reported in *Heliconius* (Van Belleghem et al. 2018), we tested whether restricting the analysis to proteins encoded by the Z-chromosome would lead to a better separation between the positive and negative controls. Remarkably, the same measures (*F*_*ST*_ and *E*_*GF*_) calculated on Z-linked genes reveal a prominent break in the continuity of the distribution (Fig. 1A). *F*_*ST*_ and *E*_*GF*_ together definitively partition all 25 pairs into two groups. Negative controls are characterized with *F*_*ST*_ < 0.14 and *E*_*GF*_ > 10%. All others (15 pairs, all positive controls and putative species pairs) form another group with *F*_*ST*_ > 0.34 and *E*_*GF*_ < 2.5%. Studies of human populations show that maximal *F*_*ST*_ between human populations is about 0.2 (Nelis et al. 2009) and humans have about 1.5–2.1% genes introgressed from Neanderthals (Wall and Yoshihara Caldeira Brandt 2016). Therefore, our criteria for species boundaries are consistent with observations made on human. The natural break in both *F*_*ST*_ and *E*_*GF*_ suggests all of the putative species (questionable species, and taxa treated as subspecies) behave similarly to the positive controls (well-defined species). Putative species are well differentiated from each other in their Z-chromosome genes, judging from high *F*_*ST*_ values and possess a well-established (but not absolute) reproductive barrier similar to the positive controls. Therefore, it is reasonable to treat these pairs of taxa as distinct species, and they are used among the different species pairs in the rest of this work.

A region of the cytochrome C oxidase subunit I (COI) encoded by the mitochondrial genome has been routinely used as a “barcode” to identify species, and sequence divergence larger than 2% in the COI barcode has been suggested as a cutoff to differentiate species (Hebert et al. 2003). We tested the ability of the COI barcode to distinguish the pairs of different species from conspecific pairs. Indeed, divergence in the barcode (x-axis in Fig. 2B) separates the true species pairs from the negative controls better than the overall average divergence of nuclear genes (y-axis in Fig. 2B). In addition, any pair with a barcode divergence larger than 2% are indeed different species. However, 8 out of the 15 pairs of different species show barcode divergence below 2%, suggesting a poor sensitivity of barcodes and calling for a need to use nuclear genomes in defining species boundaries, identifying new species, and investigating biodiversity.

### Molecular mechanisms of speciation

Genomic analysis of incipient species shows that divergence between them is frequently concentrated in certain genomic regions, termed “genomic islands of speciation” (Takuno et al. 2019). Similarly, we set out to identify genes that have diverged rapidly between species but retained relatively low polymorphism within species, and consider them to be putative “speciation genes”. We identified such genes in pairs of different species and found that they constitute about 2% ∼ 10% of all genes in each pair. We hypothesize that these putative “speciation genes” may be enriched in genes encoding phenotype divergence and reproductive barrier establishment. We mapped these proteins to Flybase (Garapati et al. 2019), so that we can compare them across different pairs. Every species pair shows higher than random overlap in putative “speciation genes” with every other species pair, and this overlap is statistically significant (p < 0.05, supplemental Table S2) for 88% of cases.

We find 244 genes that tend to (p < 0.05) be recurrently detected as putative speciation genes in multiple pairs of species (Table S3), and the most prominent recurring speciation genes are shown in Fig.3B. Gene *per*, that encode protein PERIOD, a key component of the circadian clock system, is among speciation genes for 12 out of the 15 species pairs. We observed that the most prominent speciation genes tend to have similarity in function. For example, 6 out of the 12 these genes (Fig.3B) are *trans-*regulatory elements and encode transcription factors, 8 of them encode proteins that interact with DNA or RNA, and 3 belong to circadian clock system. We further analyzed the functional enrichment (Table S4) of all 244 recurring speciation genes using Gene Ontology (GO) terms and clustered these GO terms by similarity of their meanings (Fig. 3A). This analysis reveals that divergence in circadian clock systems, protein-nucleotide interactions, immune responses, and chemical sensing may be common molecular mechanisms for speciation.

**Fig. 3.**
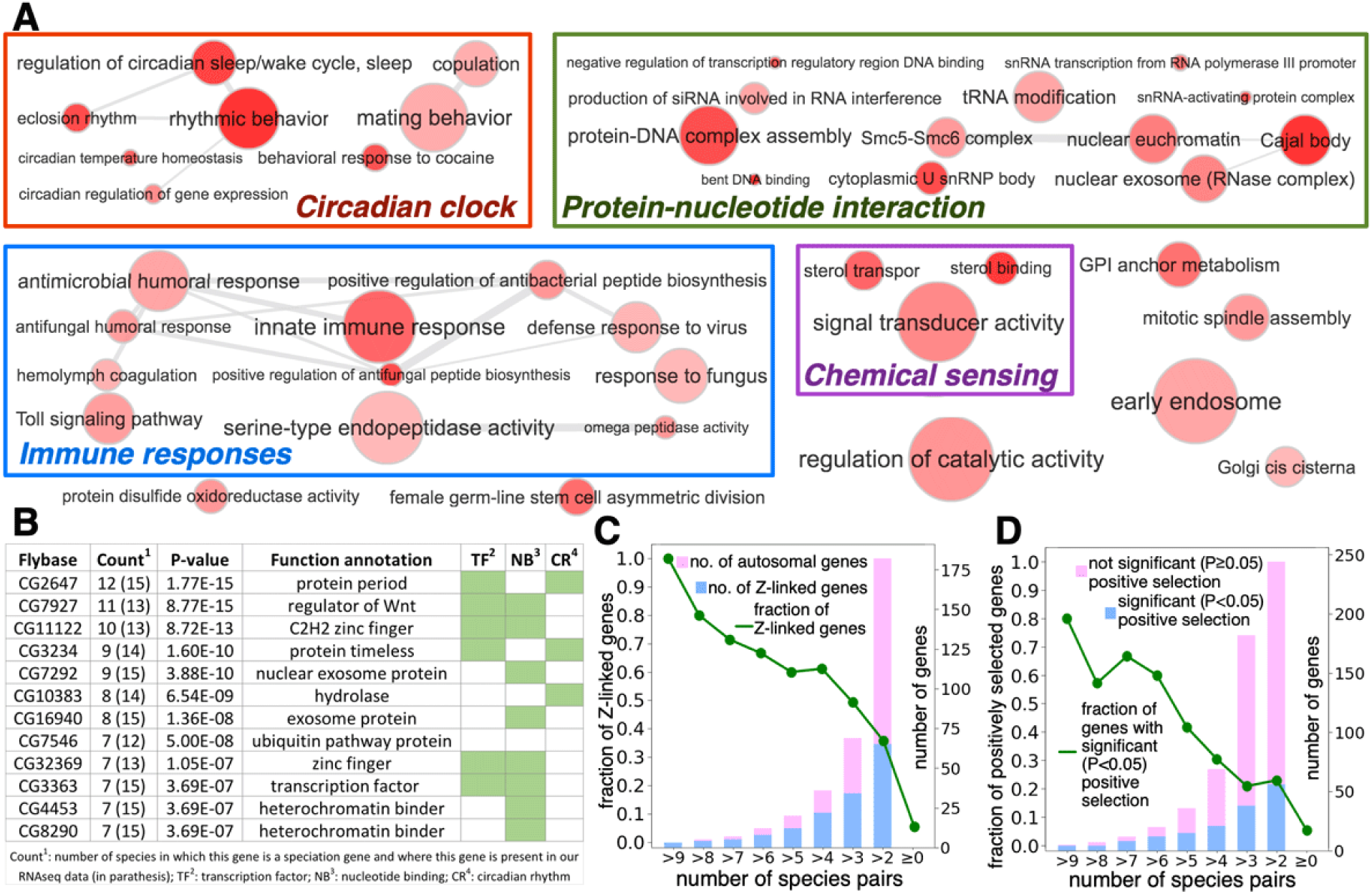
Common molecular mechanisms of speciation. (A) Functional enrichment of the recurring speciation genes reflected by GO terms. (B) Speciation genes recurring in the largest number of different species pairs. P-values indicate the chance of observing a protein at this frequency if no common speciation mechanisms are assumed. The last three columns in the table indicate whether this protein is a transcription factor, nucleotide-binding protein, or circadian clock protein, respectively. (C) Fraction (green curve) and number (blue portion of the bars) of Z-linked genes among the recurring speciation genes. (D) Fraction (green curve) and number (blue portion of the bars) of the recurring speciation genes that are positively selected.

Butterflies use chemicals, namely pheromones, to attract and select mates (Pinzari et al. 2018). Some of the recurring speciation genes encode sterol binding and sterol transporting proteins that may bind and transport pheromone molecules and thus directly mediate prezygotic isolation between species. Divergence in immune responses may be an adaptive response to exposure to different pathogens during the time of allopatric speciation, but this divergence may also play an active role in prezygotic isolation. In *Drosophila*, males inject sex peptides into females during mating, which activates the immune system of females (Hollis et al. 2019; Peng et al. 2005), protects them from possible immunogenic attacks, and thus enhances their chance to survive and produce offspring. Mating between individuals of different species may not activate the immune system to the same extent as individuals from the same species due to a lower compatibility between sex peptides and innate immune receptors from different sources. This incompatibility may reduce the fitness of females mated with males of a different species.

The importance of protein-nucleotide interaction in speciation may arise from postzygotic isolation due to Dobzhansky-Muller hybrid incompatibility (Wu and Ting 2004). Different components of the same protein-protein or protein-nucleotide complex co-evolve within each species to maintain the structure and function of the complex. If two populations have spent significant time in geographic isolation, different components of the same complex evolve independently and may lose the ability to productively interact and function. This loss could lead to hybrid incompatibility and thus contribute to the reproductive barrier. One limitation of this study based on transcriptomes is that we do not explicitly analyze *cis*-regulatory (non-coding) elements, while diversification in transcriptional regulation has been shown to play a vital role in *Drosophila* speciation (Mack and Nachman 2017; Orr 2005). However, due to the co-evolution between transcription regulatory regions and transcription factors, divergence in *cis*-regulatory elements during speciation is likely accompanied by divergence in transcription factors (i.e. proteins encoded by *trans*-regulatory elements) targeting these *cis* elements. Indeed, transcription factors, and other proteins regulating translation stand out as recurrent players in speciation, which is consistent with the findings in *Drosophila*.

In addition, recurring speciation genes show a strong tendency (P-value: 3.08e-35) to be in the Z-chromosome, and the most frequently observed speciation genes are all Z-linked (Fig. 3C). This result is consistent with the observation that *F*_*ST*_ and *E*_*GF*_ measured on Z-linked genes serve as better indicators of the speciation process than those computed on all genes. The importance of the Z-chromosome in speciation is discussed in more detail below. Finally, divergence of speciation genes between species is at least partly driven by positive selection. We identified positively selected genes using McDonald– Kreitman (MK) tests (Table S5) (McDonald and Kreitman 1991), which detects genes showing elevated nonsynonymous substitution rates between species compared to within species. A significant portion of recurring speciation genes (23%) are positively selected (P < 0.05), much larger (P-value: 8.1e-21) than the fraction of positively selected genes in the entire transcriptome (5.3%, Fig. 3D). This observation is again consistent with studies in other species, suggesting that speciation genes are typically fast evolving and positively selected (Orr 2005).

### Circadian clock proteins in speciation

All four key components of the circadian clock system (Fig. 4A), CLOCK (CLO), CYCLE (CYC), PERIOD (PER), and TIMELESS (TIM), are encoded by the recurring speciation genes we identified, with PERIOD being the most prominent one. Expression of PER and TIM are activated by CLO and CYC, and accumulation of PER and TIM lead to formation of the PER/TIM complex, which is then transported to the nucleus and suppresses the transcription factor activity of CLO and CYC (Young 1998). Circadian clock proteins are suggested to be directly involved in courtship behavior: they control the daily rhythm of mating behavior (Sakai and Ishida 2001) and the species-specific mating song (Emmons and Lipton 2003) in *Drosophila*.

**Fig. 4.**
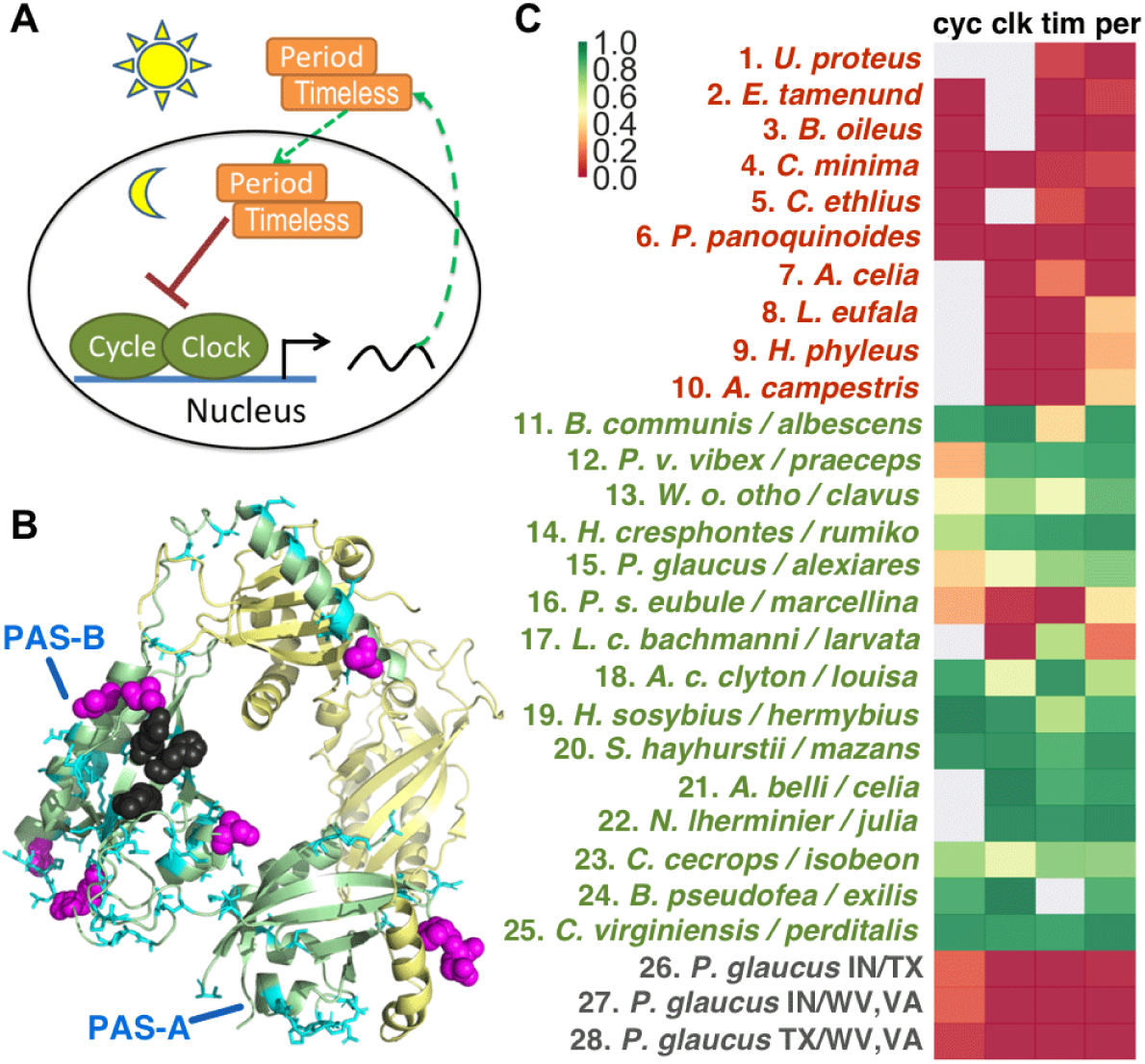
Circadian clock proteins play important roles in speciation. (A) a schematic of the circadian clock system. (B) Variations in protein PERIOD between species pairs mapped to the 3D structure (PDB id: 3RTY). The structure includes the two PAS domains of PER: PAS-A and PAS-B. Positions that differ in one pair of species are shown as cyan sticks and those that differ in more than one pair are shown as magenta spheres. Residues that disrupt the interaction between PER and TIM are shown as black spheres. (C) *F*_*ST*_ (red - low, green - high) of the 4 circadian clock proteins for 25 pairs of counterparts across central Texas suture zone and for populations of *Pterourus glaucus* at different latitudes. Genes whose transcripts are absent in the RNA-seq dataset are shown as gray squares.

We mapped the variations (shown as cyan sticks in Fig. 4B) occurring in each pair of species onto the 3D structure of *Drosophila* PER (King et al. 2011), and identified positions (magenta spheres in Fig. 4B) with recurring variations. They concentrate in the second PAS domain of PER, which interacts with TIM (Hennig et al. 2009). Positions suggested to be crucial for the PER/TIM interaction in mutagenesis experiments (Hennig et al. 2009) are shown as black spheres in Fig. 4B, and several recurring variations are adjacent to these positions. These variations may modulate the binding affinity between PER and TIM, altering the circadian rhythms of different species and contributing to their adaptation to different climate and photoperiod. Due to the important role of circadian clock proteins in mating rhythm (Allada and Chung 2010; Merlin et al. 2007), such divergence in clock proteins may make different species incompatible in their preferred mating time and contribute to prezygotic isolation.

Furthermore, divergence in circadian clock genes can lead to postzygotic isolation due to Dobzhansky-Muller hybrid incompatibility (Wu and Ting 2004). PER and TIM of the same species coevolve to maintain proper affinity for triggering the formation of the PER/TIM heterodimer at the right time, and to ensure a period of about 24 hours. This 24-hour cycle accommodates crucial daily activities such as feeding and mating. However, in a hybrid individual with PER and TIM from different species that have not been evolving together, dimerization between PER and TIM may be affected, impairing the 24-hour cycle and reducing the fitness of the hybrid.

Contribution of clock proteins to both prezygotic and postzygotic isolation make them good markers for speciation. We computed *F*_*ST*_ for circadian clock genes and for each of the 25 pairs of counterparts across the central Texas suture zone (Fig. 4C), and observed a much higher *F*_*ST*_ for pairs that are different species than conspecific pairs, with only a few exceptions such TIM and CLO in *Phoebis*, and CLO and PER in *Libytheana.* It is important to note that divergence in circadian clock is not a necessary outcome of population separating into different latitudes. For example, we computed the *F*_*ST*_ for populations of *Pterourus glaucus* in different latitudes and geographic regions (Indiana, Virginia-West Virginia, and Texas, separated by 1000 miles). These populations do not show divergence in circadian clock genes (last three rows, #26-28 in Fig. 4C), despite signs of adaptation to latitudes as reflected by the difference in the number of generations per year (Cong et al. 2015b). In contrast, the species pair *P. glaucus* and *P. alexiares* (#15 in Fig. 4C) reveals a notable difference in the clock proteins despite smaller distance between these populations (300 miles).

### Introgression and the role of Z-chromosome in speciation

Butterflies, like birds, but in contrast to mammals and *Drosophila*, use the ZW sex determination system (Traut et al. 2007), where a female has two dissimilar chromosomes (ZW) and a male has a pair of similar chromosomes (ZZ). The importance of the Z-chromosome in speciation revealed in our analysis is not due to a larger divergence of the Z-chromosome between species compared to autosomes: the divergence in the Z-chromosome is slightly lower than in autosomes (Fig. 5A, left). Instead, the superior performance of the Z-chromosome in spotting speciation events is probably because Z-chromosome genes have a lower chance of introgression between species (Fig. 5A, middle). Reduced introgression in sex chromosomes have been observed in *Heliconius* butterflies used as model organism for introgression studies (Martin et al. 2013) and in human (Sankararaman et al. 2014). Gene flow between diverged species increases intraspecific divergence and lowers interspecific divergence. Therefore, elevated introgression in autosomes compared to the Z-chromosome results in a lower *F*_*ST*_ in autosomes (Fig. 5A, right). Reduced introgression from Neanderthal in human X-chromosome compared to autosomes was suggested to be related to sex-biased hybridization (Sankararaman et al. 2014; Wolf and Akey 2018), where interbreeding mostly happened between Neanderthal males and modern human females. A male Neanderthal had only 50% chance to bring its X-chromosome into the gene pool of *Homo sapiens*. In addition, Neanderthal X-linked genes in modern human genetic background may decrease fertility in males (Sankararaman et al. 2014), thus explaining the lower frequency of X-linked introgression than autosomal introgression in modern human genomes.

**Fig. 5.**
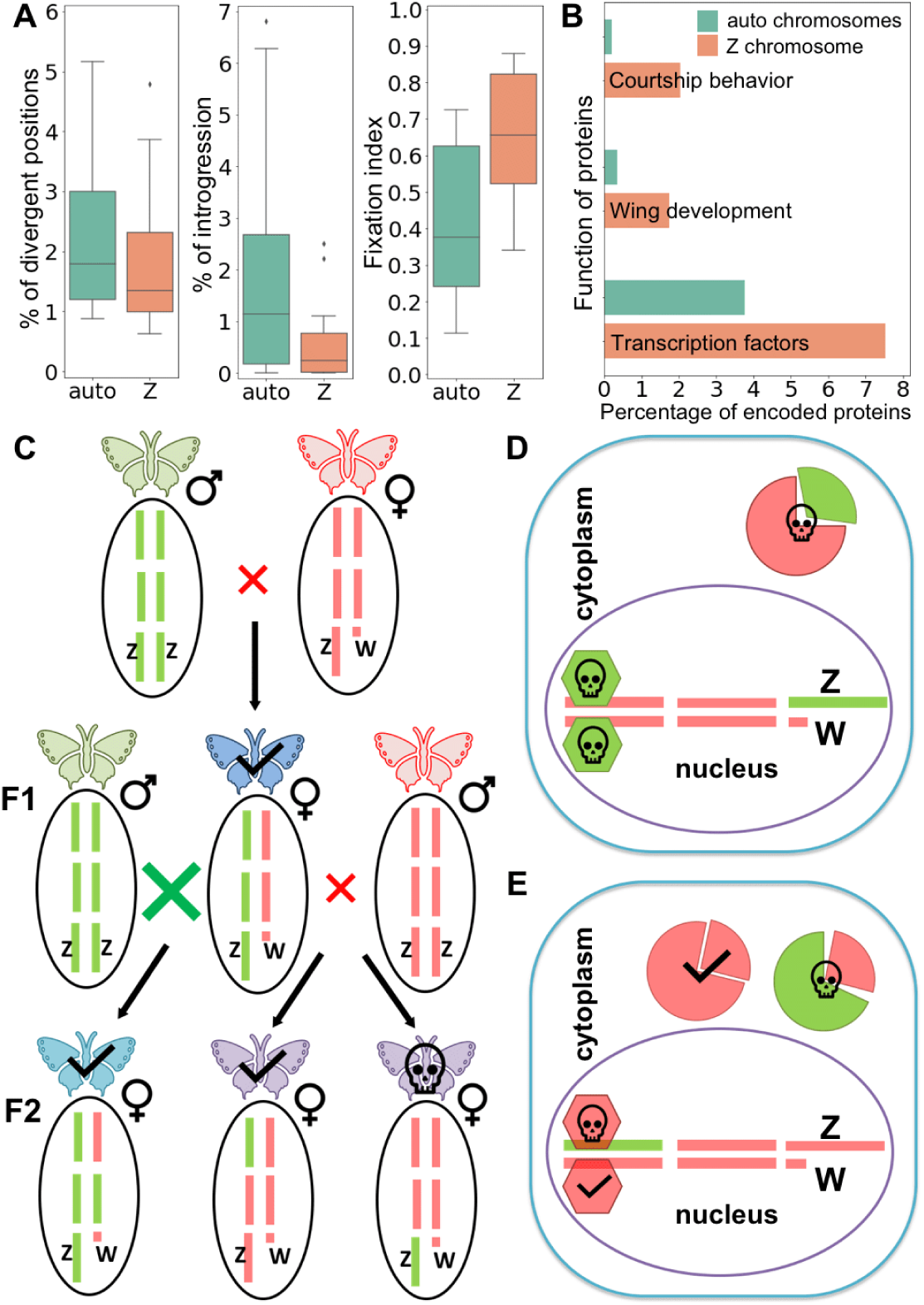
Z chromosomes of butterflies are resistant to introgression. (A) The level of divergence, introgression and fixation indices for 15 pairs of species computed on autosomes and Z-chromosomes. (B) Z-chromosome encodes higher fraction of proteins related to courtship behavior, wing development and transcription factors than autosomes. (C) A model of chromosome introgression explains why Z-chromosome is resistant to introgression. Each individual is marked by a butterfly icon and an oval under the icon displays its genotype. Green bars marks chromosomes from the green butterfly, and red bars mark those from the red one. Crosses indicate mating and a larger cross indicates a higher chance to mate. A skull on a butterfly indicates significant reduced fitness or death, while a check indicates a higher chance of survival. (D) and (E): The effect of Dobzhansky-Muller hybrid incompatibility is stronger in cases of (D) Z-chromosome than (E) autosome introgression. Bars inside the purple circle are chromosomes and hexagons represent transcription factors encoded by Z chromosomes. Cytoplasmic proteins encoded by Z chromosomes are shown as minor sectors while those encoded by autosomes are shown as major sectors. The color of these shapes indicates the origin of these molecules.

Analysis of Z-linked gene function in butterflies revealed significant enrichment in proteins participating in courtship, wing development, and transcription factors controlling important developmental processes (Fig. 5B). Additionally, the Z-chromosome is enriched in genes involved in morphogenesis, development of neural systems, and signaling pathways mediated by hormones (Table S6). Such enrichment suggests a prominent role of the Z-chromosome in determining traits that are subject to sexual selection and in promoting copulation between individuals with similar Z-chromosomes. For example, 3 of the 4 circadian clock genes are encoded by the Z-chromosome in *Heliconius* (Heliconius Genome Consortium 2012), and thus the Z-chromosome is more important in determining the rhythm of mating. As a result, we propose the following model to explain the Z-chromosome’s resistance to introgression and importance in speciation in butterflies (Fig. 5C).

When two different species (male A and female B) meet, due to the porous reproductive barrier (Mallet et al. 2016), they occasionally mate and produce F1 hybrid offspring. The F1 hybrids are likely to have lower fitness, and thus males among them are less likely to be favored by sexual selection. However, since females dominate sexual selection in butterflies, the F1 females still have a chance to mate and pass introgressed genes to their offspring. F1 female gets the Z chromosome from species A (father), and thus it will have a higher chance to mate with species A due to similar sexually selected traits encoded by their Z-chromosomes. Mating between F1 hybrid female and species A leads to introgression of autosomal genes, but not Z-linked genes from species B to species A.

If the F1 hybrid female mates with a male from species B, it will produce F2 offspring with introgressed autosomes and/or Z chromosome. However, the F2 females with Z chromosome introgression are expected to have much lower fitness due to incompatibility between Z chromosome and autosomes of a different species as illustrated in Fig. 5D. This incompatibility is reflected both in protein-protein complexes and protein-DNA complexes. Since the single-copy Z chromosome encodes a number of important transcription factors (TF), the incompatibility between these TFs and the *cis-*regulatory elements from another species will likely have a decisive impact on the survival or fitness of the F2 females. In contrast, individuals with introgressed autosomes will not suffer as much from the incompatibility of molecules from different origins, because the non-introgressed homologous chromosome of the introgressed autosome can encode proteins that are compatible with the genetic background of the F2 hybrid (Fig. 5E).

This model suggests that in an organism with ZW sex determination, an introgressed Z chromosome has a much lower chance to spread in the population through females. Although males can spread introgressed Z chromosomes, their role in driving the population’s gene pool is much weaker because almost every female has the chance of producing offspring, while males compete for the chance of mating and only a fraction succeed. Homologous recombination breaks the linkage of an introgressed segment or chromosome, allowing shorter pieces of foreign Z-chromosomes to spread in the population, but it does not change the fact that on average, introgressed Z-linked segments may reduce fitness more than autosomes. Occasionally, some introgressed alleles may increase the fitness, and they can spread throughout the population and even completely replace the endogenous alleles due to selective sweeps (Messer and Petrov 2013). Due to the reduced chance of introgression, the Z chromosome more readily carries the history of a species than autosomes and, contrary to the mitogenome, does not readily introgress into a different species. Even more, the important role of Z-linked genes in morphogenesis and courtship may promote the introgression of genes on other chromosomes: as long as a hybrid female has the Z-chromosome of one species, it will have a high chance to spread its genes in this species. High level of introgression may be beneficial for a species, despite the expected cost in fitness to an individual. Even if most of the foreign genes are not compatible with the genetic background of a species, some can bring a selective advantage and allow a species to adapt quickly to different environments, foodplants, or pathogens.

## MATERIALS AND METHODS

We focused on populations around the central Texas suture zone and assembled 25 pairs of counterparts from both the eastern and western side of the suture zone. For each pair, we gathered from 2 to 15 specimens on each side of the suture zone. A minimum of 2 specimens on either side of are needed to estimate the variability within each population. The information about the specimens is in Table S1. For samples from the genus *Calycopis*, we used genomic DNA data we published before (Cong et al. 2017; Cong et al. 2016a). For samples from the genera *Phoebis* and *Pterourus*, we used genomic DNA libraries (Cong et al. 2015b; Cong et al. 2016b). For all other genera, we obtained RNA-seq libraries.

### Library preparation and sequencing

Specimens for RNA-seq libraries were euthanized upon capture by thorax pinching, and the whole specimen bodies except wings and genitalia were preserved in *RNAlater* solution. Total RNA was extracted from each specimen using QIAGEN RNeasy Plus Mini Kit, and mRNA was further isolated using NEBNext Poly(A) mRNA Magnetic Isolation Module. RNA-seq libraries were prepared with NEBNext Ultra Directional RNA Library Prep Kit following manufacturer’s protocol. We pooled RNA-seq libraries of every 12 specimens together and sequenced each pool using one illumina Hiseq2500 lane for 150 bp at each end. Specimens for genomic DNA libraries were either collected in the field (and stored in *RNAlater* or EtOH) or borrowed from collections listed in the Acknowledgements. A piece of thoracic tissue from fresh specimens, and either the abdomen or a leg from pinned museum specimens were used for DNA extraction with Macherey-Nagel NucleoSpin tissue kit following the manufacturer’s protocol. Genomic DNA libraries was made using NEBNext® Ultra(tm) II DNA Library Prep Kit. We pooled DNA libraries according to the expected size of their genome and typically targeted 10X coverage for each sample. We sequenced the pooled genomic libraries using Hiseq X ten sequencing service from Genewiz. The sequence reads will be deposited to the NCBI SRA database upon publication.

### Assembling reference transcriptomes for each case

After removal of contamination from sequencing adapters and the low-quality portion (quality score < 20) using AdapterRemoval (Schubert et al. 2016), we applied Trinity (version r20140413p1) (Haas et al. 2013) to *de novo* assemble the transcriptome separately for each specimen. For each of the 25 cases, we merged the transcripts from all specimens in each pair (they are all closely related) and mapped them to the protein set of a reference genome. We used 6 reference genomes (and the mitogenomes of these species) including *Cecropterus* [formely Achalarus] *lyciades* (Shen et al. 2017), *Calycopis cecrops* (Cong et al. 2016a), *Danaus plexippus* (Zhan et al. 2011), *Heliconius melpomene* (Heliconius Genome Consortium 2012), *Lerema accius*, (Cong et al. 2015a) and *Pterourus glaucus* (Cong et al. 2015b), and for each case, the most closely related reference was used. We applied BLASTX (e-value: 0.00001) (version 2.2.31+) (Altschul et al. 1997) to map the transcripts to proteins. Transcripts that could not find a confident (e-value <= 0.00001) hit among reference proteins were discarded. We filtered the BLASTX hits requiring the aligned positions between the transcript and the hit to cover at least 50% of the residues in the hit, and the remaining hits were ranked primarily by e-value and secondarily by bit score. From the ranked list we identified the best hits that were aligned to non-overlapping regions in the transcript. Usually there was only one best hit, and in cases where multiple non-overlapping best hits were identified, the transcript was split to multiple segments corresponding to multiple best hits.

Each transcript was considered to map to the top hit from the reference protein set. Transcripts mapping to the same reference protein were aligned against each other using BLASTN (version 2.2.31+) (Altschul et al. 1997) to remove redundancy and to merge partial transcripts to a complete one. We wanted to represent alternatively spliced isoforms with just the longest isoform, and we removed other isoforms and redundant transcripts from different specimens using the following criteria: (1) if two transcripts were over 95% identical to each other and the aligned region covered at least 80% of one transcript, the shorter transcript was removed; (2) if two transcripts were over 90% identical to each other, their aligned region covered at least 80% of one transcript, and the two transcripts share at least one identical 40mer, the shorter transcript was removed. In order to merge the partial transcripts, we referred to the alignment between transcripts and proteins from reference genomes. If two transcripts were aligned to different portion of the protein with at least 20 residues as overlap, the two transcripts were merged and in the overlapping region, the sequence more similar to the reference protein was taken.

### Obtaining sequence alignments

For specimens of *Pterourus, Calycopis*, and *Phoebis*, we sequenced the DNA libraries since the reference genomes for these genera were available. We aligned the genomic DNA reads to the reference genomes using BWA (version 0.6.2-r126) (Li and Durbin 2009) and performed SNP calling with GATK (version 3.3-0) (DePristo et al. 2011). The sequence of each specimen was derived from the GATK results, and the transcript sequences was extracted from these genomic sequences. For specimens of all other genera, where we used RNA-seq reads, a similar pipeline of BWA and GATK was applied using the assembled transcriptomes as references. Transcripts that were covered for at least 60bp by at least two samples from the west and two from the east of the suture zones were kept, resulting in 10675 - 16494 transcripts in each case.

### Detecting Z-linked transcripts

High conservation of gene content has been reported in Lepidoptera Z chromosome (Fraisse et al. 2017), and therefore we can deduce the Z-linked genes in other species by comparing to *Heliconius melpomene* reference genome, where Z chromosome sequence was known (Heliconius Genome Consortium 2012). We identified the Z-linked transcripts in each of the 25 cases using a reciprocal best hit approach. We used the BLASTX to search each query transcript against all the *Heliconius* proteins to identify the most closely related hit. If the top hit was a Z-linked protein, we examined whether the hit also found the query transcript as the closest hit by TBLASTN search against all the transcripts from this case. If the answer was yes, we assigned the query transcript as Z-linked.

### Designing measurements to separate species from non-species pairs

For each of the 25 pairs, we computed the fixation indices (*F*_*ST*_) and extent of gene flow (*E*_*GF*_). We computed these measures both for all the transcripts, and for Z-linked transcripts only. The 25 pairs consist of both different species and conspecific populations, but we use the term “populations” for all pairs below for simplicity. *F*_*ST*_ is calculated as 1-DIV_within_/DIV_between_, where DIV_within_ and DIV_between_ are the average divergence between a pair of specimens within the same population and between different populations, respectively.

If one allele in population A was more similar to alleles from population B than alleles from its own population, we considered it to possibly result from gene flow from population B to A. We split each transcript into windows, and we varied the window size from 1.0 kbp to 2.0 kbp. We examined one sample in each window at a time and attempted to assemble a putative foreign allele that is the most similar to the other population by selecting SNPs that are more similar to the other population at each position. We obtained the divergence (*div1*) between this putative foreign allele and the other population, as well as its divergence (*div2*) from its own population. In addition, we computed the divergence (*div3*) between the two populations using other samples excluding the one being examined. We considered a putative foreign allele to be an “admixed fragment” if *div2* - *div1* > *div3*. The fraction of admixed fragments in each sample was then calculated as the number of admixed fragments divided by twice the total number of windows. We multiplied the total number of windows by two because butterfly genomes are diploid, and we only detected a haplotype as admixed for each sample in each window. We averaged the fraction of admixed fragments obtained from different window size and in different samples to obtain a final estimate of the extent of gene flow (*E*_*GF*_) between the two populations.

### Identification of putative speciation genes

Our analysis revealed 15 pairs of counterparts across the central Texas suture zone as species pairs, and we further identified the putative speciation genes for each pair. We hypothesized that the proteins showing significant divergence between species but are relatively conserved within each species are more likely to be encoded by speciation genes, and we identified them using protein sequences by two criteria. First, we required the *F*_*ST*_ for this protein to be among the top 20% of all proteins, and this criterion ensured that the selected proteins were relatively conserved within a species. Second, we detected all the positions that were conserved (sharing a common amino acid in over 75% of sequences) within but different between species, and further identified proteins that were significantly enriched (p < 0.05) in such positions. The enrichment was quantified using binomial tests (p = rate of divergent positions in the alignments for all proteins, m = the number of divergent positions in this protein, N = the total number of aligned positions in this protein).

In order to identify the common speciation genes, we mapped all the reference transcripts to *Drosophila* proteins in Flybase (Garapati et al. 2019) by identifying the top BLASTX hits. We counted how many times each Flybase entry appeared as putative speciation genes among all 15 species pairs. If multiple (n) transcripts in one case were mapped to the same Flybase entry, each transcript was counted as 1/n of that entry. Some genes tend to be among speciation genes for multiple cases: when this tendency was significantly larger than random, we considered this gene to be a recurring speciation gene. We simulated random process of selecting x genes in each case, respectively, where x was the actual number of putative speciation genes we identified in each case. We repeated the random simulation 10,000 times, and for each gene, we selected 5% of simulations showing the highest frequency for this gene. The lowest frequency of observing the gene among the selected simulations for this gene was used as to a cutoff for an observed frequency to be significantly higher than random (P-value < 0.05).

### Detecting positively selected genes between species

We tested whether a gene was positively selected during the divergence of 15 pairs of butterfly species using McDonald–Kreitman (MK) test (McDonald and Kreitman 1991). For each species and each gene, we estimated the number of nonsynonymous substitutions (P_N_) and synonymous substitutions (P_S_) needed to change from the codons of one individual (A) to the codons of another individual (B). If there were multiple substitution paths to change from the codon of A to the codon of B, the path with the smallest number of nonsynonymous substitutions (parsimonious) was taken. For each gene, P_N_ and P_S_ values for all 30 species in the 15 pairs were summed to get total P_N_ (TP_N_) and P_S_ (TP_S_). Similarly, for each pair of species and each gene, we counted the number of nonsynonymous substitutions (D_N_) and synonymous substitutions (D_S_) to change from the codons of one species to the codons of another species. The total D_S_ (TD_S_) and total D_N_ (TD_N_) were summed over 15 species pairs for each gene. To calculate the statistical significance for positive selection in each gene, we used Fisher’s exact test to compare if TD_N_/TD_S_ was significantly larger than TP_N_/TP_S_. A gene with P-value less than 0.05 was considered to be positively selected.

### Functional enrichment analysis

We identified the enriched GO terms associated common speciation genes using binomial tests (m = the number of common speciation genes that were associated with this GO term, N = number of common speciation genes, p = the probability for this GO term to be associated with any gene). GO terms with P-values lower than 0.05 were considered enriched, and those with P-values less than 0.03 were shown in Fig. 3A. A similar analysis was performed on Z-linked genes.

## ACKNOWLEDGMENTS

We acknowledge Texas Parks and Wildlife Department (Natural Resources Program Director David H. Riskind) for the permit #08-02Rev that makes research based on material collected in Texas State Parks possible; Paul A. Opler and Boris Kondratieff (Colorado State University Collection, Fort Collins, CO, USA), Jerry A. Powell (Essig Museum of Entomology, University of California, Berkeley, CA, USA), Edward G. Riley, Karen Wright, and John Oswald (Texas A & M University, College Station, TX, USA), and Robert K. Robbins, John M. Burns, and Brian Harris (National Museum of Natural History, Smithsonian Institution, Washington, DC, USA) for granting access to the collections under their care and for stimulating discussions. The study has been supported by grants from the National Institutes of Health GM094575 and GM127390, and the Welch Foundation I-1505 to NVG.

