## Supplementary figures and images for "Genomic determinants of speciation"

### Figs. 1 and 2

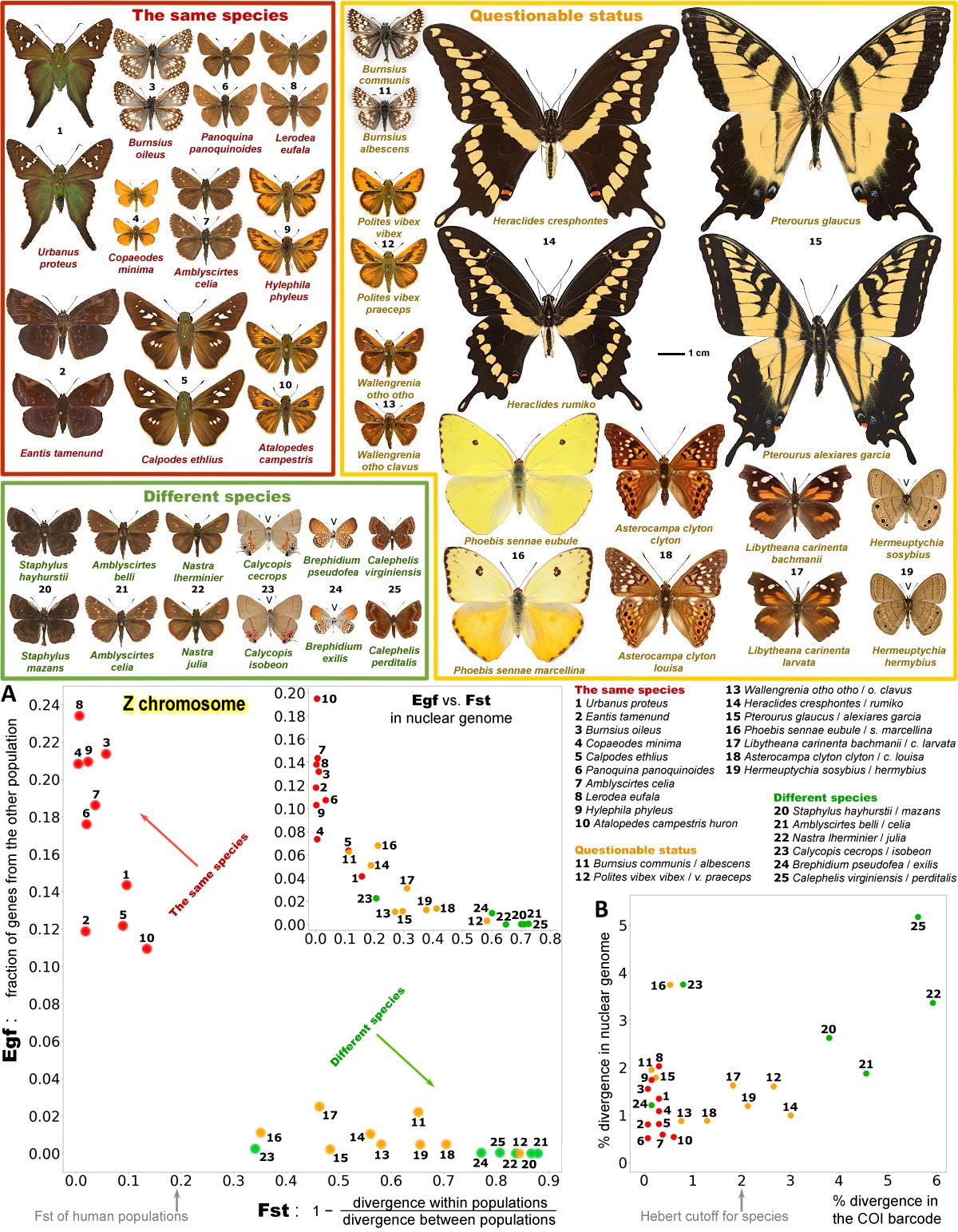
